# Longitudinal investigation of changes in resting-state co-activation patterns and their predictive ability in the zQ175 DN mouse model of Huntington’s disease

**DOI:** 10.1101/2022.10.09.511485

**Authors:** Mohit H. Adhikari, Tamara Vasilkovska, Roger Cachope, Haiying Tang, Longbin Liu, Georgios A. Keliris, Ignacio Munoz-Sanjuan, Dorian Pustina, Annemie Van der Linden, Marleen Verhoye

**Affiliations:** Bio-Imaging Lab, University of Antwerp, Antwerp, Belgium; µNEURO Research Centre of Excellence, University of Antwerp, Antwerp, Belgium; CHDI Management/CHDI Foundation, Princeton, NJ, United States of America; Institute of Computer Science, Foundation for Research & Technology – Hellas, Heraklion, Crete, Greece

**Keywords:** Huntington’s disease, Animal models, Resting-state, functional MRI, Co-activation patterns, Classification, machine-learning

## Abstract

Huntington’s disease (HD) is a neurodegenerative disorder caused by expanded (≥40) glutamine-encoding CAG repeats in the huntingtin gene, which leads to dysfunction and death of predominantly striatal and cortical neurons. While the genetic profile and behavioural signs of the disease are better known, changes in the functional architecture of the brain, especially before the behavioural symptoms become apparent, are not fully and consistently characterized. In this study, we sought markers at pre, early and late manifest states of phenotypic progression in the heterozygous (HET) zQ175 delta-neo (DN) mouse model, using resting-state functional magnetic resonance imaging (RS-fMRI). This mouse model shows molecular, cellular and circuitry alterations that resemble those seen in HD in humans. Specifically, we investigated, longitudinally, changes in co-activation patterns (CAPs) that are the transient states of brain activity constituting the resting-state networks (RSNs). Most robust changes in the temporal properties of CAPs occurred at the late manifest state; the durations of two anti-correlated CAPs, characterized by simultaneous co-activation of default-mode like network (DMLN) and co-deactivation of lateral-cortical network (LCN) and vice-versa, were reduced in the zQ175 DN HET animals compared to the wild-type mice. Changes in the spatial properties, measured in terms of activation levels of different brain regions, during CAPs were found at all three states and became progressively more pronounced at the manifest states. We then assessed the cross-validated predictive power of CAP metrics to distinguish HET animals from controls. Spatial properties of CAPs performed significantly better than the chance level at all three states with 80% classification accuracy at the early and late manifest states.

## Introduction

Huntington’s disease is an autosomal dominant, neurodegenerative disorder that manifests itself typically in adulthood as a combination of motor and cognitive symptoms that worsen progressively (Bates et al., 2015). Neuropsychiatric symptoms such as depressed mood, mania, anxiety, obsessions-compulsions, and psychosis have also been reported in HD patients (Paoli et al., 2017). HD is caused by expanded (≥40) glutamine-encoding CAG repeats in the huntingtin gene that lead to the expression of mutant huntingtin (mHTT) (Ross and Tabrizi, 2011). Brain regional atrophy starts in the striatum and as the disease progresses the degeneration expands to cortical regions (Nanetti et al., 2018; Rosas et al., 2008). As in case of other neurodegenerative diseases such as Alzheimer’s disease (AD) and Parkinson’s disease, neurodegeneration in HD can start before the appearance of clear behavioral signs. Neuroimaging investigations can contribute greatly in identifying disease markers before making a clinical motor diagnosis. Resting-state functional magnetic resonance imaging (RS-fMRI), in particular, has been a promising tool in developing such biomarkers for a number of neurological as well as neurodegenerative disorders such as stroke (Baldassarre et al., 2014; Siegel et al., 2016), disorders of consciousness (Di Perri et al., 2018) and AD (Badhwar et al., 2017). In AD, alterations in the functional connectivity (FC) of the default mode network (DMN), the most prominent of the resting-state networks (RSNs), have been shown to correlate with amyloid-β deposits (Buckner et al., 2005). In the case of HD, alterations in the RS-FC have been found in several RSNs including visual, somato-motor, executive, auditory and cerebellar networks (Poudel et al., 2014; Werner et al., 2014; Wolf et al., 2014), mostly during the manifest stage of the disease. However, the findings have not been consistent in regards to the directionality of the change in patients vis-à-vis healthy participants (Pini et al., 2020).

Most investigations of RS-FC in HD have relied on the methods of static analysis of RS-fMRI data, namely the seed-based analysis, independent component analysis or graph-theoretical analysis on the static FC (Pini et al., 2020). The assumption underlying these analysis methods is that an estimate of FC derived from the entire duration of a RS scan is a robust representation of the subject’s brain state. This assumption, however, ignores temporal fluctuations of FC over the duration of the scan that reveal the brain dynamics at shorter time-scales constituting the FC and RSNs. In the last decade, novel techniques to extract dynamic changes in the RS data during a scan have been developed (Allen et al., 2014; Deco et al., 2017; Hutchison et al., 2013). One stream in this emerging research domain has focused on quantifying variations within the static FC and testing the non-stationarity of FC with rigorous methods (Hindriks et al., 2016; Zalesky et al., 2014). Another stream investigates the topology and dynamics of transient brain-states that constitute the RSNs and their neural correlates (Liu and Duyn, 2013a; Liu et al., 2018; Majeed et al., 2011; Thompson et al., 2014). Co-activation patterns (CAPs) (Liu and Duyn, 2013a) are an example of patterns of brain activity that can be extracted from the dynamical analysis of RS-fMRI data representing blood-oxygenation-dependent (BOLD)-based transient brain states that are obtained at a single time frame resolution (i.e., at every measured fMRI volume). Another example is the quasi-periodic patterns, which are recurring spatiotemporal patterns of BOLD activity of fixed temporal length (Belloy et al., 2018a; Majeed et al., 2011). The time-varying information extracted with these methods has shown that traditional static FC analyses conceal important clues on the temporal dynamics of brain activity, such as the specific micro-states, their occurrence, their order, and their duration. Such information can provide additional features with promising value for being a sensitive biomarker in neurodegenerative diseases like HD.

Here we investigated, longitudinally, changes in the RS-CAPs that are transient constituents of RSNs in a mouse model of HD. The mouse model is the knock-in heterozygous (HET) zQ175 delta-neo (DN) that shows molecular, cellular and circuitry alterations that resemble those observed in people with HD (PwHD) (Heikkinen et al., 2020; Menalled et al., 2012; Southwell et al., 2016). In this mouse model, motor disturbances are manifested at 6-months of age and worsen at 10 months of age (Heikkinen et al., 2020). We acquired longitudinal RS-fMRI data in wild-type (WT) and HET animals at these progression states: premanifest (3 months), early manifest (6 months), and late manifest (10 months). We identified CAPs using a recently developed method (Gutierrez-Barragan et al., 2019) at each state. We hypothesized that the impact of mutant huntingtin (mHTT) will be reflected in the spatial (activation pattern) and temporal (frequency and duration) components of prominent and physiologically informative RS-CAPs at both premanifest and manifest progression states. We further hypothesized that reliable CAP metrics should be able to accurately predict the identity of HET mice and paid particular attention to their ability to classify animals in the premanifest state.

## Materials and Methods

### Ethical statement

All procedures were performed in strict accordance with the European Directive 2010/63/EU on the protection of animals used for scientific purposes. The protocols were approved by the Committee on Animal Care and Use at the University of Antwerp, Belgium (permit number 2017-09) and all efforts were made to minimize animal suffering.

### Animals

zQ175 delta neo (DN, neomycin-sensitive cassette excised), knock-in heterozygous mouse model is used in this study. The neo+ version of the model used in earlier studies showed a close resemblance in molecular, cellular and circuitry alterations of human Huntington’s disease (Heikkinen et al., 2020; Menalled et al., 2012). The phenotypic conversion in the heterozygous zQ175 DN model (Southwell et al., 2016) happens at 6-months of age, and beyond 7 months, the animal is considered to be in the manifest state. The cohort used in this study consisted of 20 male zQ175 DN heterozygous (referred henceforth as HET) mice and 19 age-matched wild-type (WT) littermates. RS-fMRI data were collected longitudinally at three time points when the animals were 3, 6, and 10 months of age. One animal from each group was found to be mis-genotyped, hence both animals were excluded from the analysis. Thus, the analysis was performed on 18 WT and 19 HET animals.

### RS-fMRI data acquisition

The mice were initially anesthetized with 2% isoflurane (IsoFlo, Abbott, Illinois, USA) in a mixture of 200 ml/min nitrogen and 400 ml/min oxygen. Heads were fixed with bite and ear bars. Ophthalmic ointment was applied to the eyes. Animal core body temperature was monitored with a rectal temperature probe and kept stable at 37 °C via hot air supply (MR-compatible Small Animal Heating System, SA Instruments, Inc.). A pressure sensitive pad and a fiber-optic pulse oximeter placed over the tail were used to monitor the breathing rate and the heart rate & O2 saturation (MR-compatible Small Animal Monitoring and Gating system, SA Instruments, Inc.) respectively. Following animal handling, the mice received a bolus injection of medetomidine (0.075 mg/kg; Domitor, Pfizer, Karlsruhe, Germany) after which the isoflurane level was reduced to 0.4% over the course of 30 min. Throughout the entire imaging protocol, the isoflurane level was maintained at this level. A subcutaneous catheter allowed continuous infusion of medetomidine (0.15 mg/kg/hr) starting 10 minutes after bolus injection, resembling a brain state of rest (Jonckers et al., 2014). The RS-fMRI acquisition started 40 minutes after bolus injection. After the MRI procedure, the effect of medetomidine was counteracted by injecting 0.1 mg/kg atipamezole (Antisedan, Pfizer, Karlsruhe, Germany).

MRI scans were performed on a 9.4 T Biospec Bruker system (Germany) using a mouse head receiver 2×2 array cryo-coil. RS-fMRI data were acquired with a single shot gradient-echo echo-planar imaging (EPI) sequence, field of view (27 × 21) mm^2^, matrix dimensions (MD) [90,70], 12 horizontal slices with 0.4 mm thickness, in-plane resolution (300×300) µm^2^, flip angle 60°, bandwidth 400 kHz, repetition time (TR) 500 ms, echo time (TE) 15 ms, 1200 repetitions. 3D RARE images were acquired, in order to create a study specific 3D template, with TR 1800 ms, TE 42 ms, MD [256,256,128], isotropic resolution (78 × 78 × 78) µm^3^.

### Pre-processing of functional MRI data

RS-fMRI data of each subject were realigned to the first time frame using the least-squares approach and a 6-parameter rigid body spatial transformation. A study-based 3D template was generated in Advanced Normalization Tools (ANTs), using individual 3D RARE images from 1/3 of the subjects from each group at each time point. Transformation parameters were extracted from two different normalization steps, where each RS-fMRI scan was registered to the corresponding subject 3D RARE image acquired at the same time point, and each subject 3D RARE scan was normalized to the study-based template. Moreover, spatial normalization parameters were calculated between the study-based template and an in-house C57BL6 mouse brain atlas. Finally, the RS-fMRI data were spatially normalized to the in-house atlas by combining the previous normalization estimation parameters (RS-fMRI to 3D RARE, 3D RARE to study template and study template to in-house atlas). Next, we performed in-plane smoothing using a Gaussian kernel with full width at half maximum of twice the voxel size. Motion vectors were then regressed out of the image-series. These procedures were performed using Statistical Parametric Mapping (SPM12) software (Welcome Department of Cognitive Neurology, London, UK).

Images were then filtered voxel-wise using a 0.01-0.2 Hz butterworth band-pass filter, and quadratic detrended. Five time frames at the start and the end of the scan were excluded before and after filtering to avoid transient effects and to minimize boundary effects of filtering respectively. Global signal, obtained using a whole brain mask, was regressed out from the BOLD signal of each voxel that was then normalized to zero mean and unit variance. We used in-house MATLAB scripts for filtering, detrending, global signal regression and z-scoring.

### CAP analysis

Since cerebellar BOLD signal was found to be anti-correlated with most regions in the cerebrum, this strong anti-correlation masked the relationships between regions of the cerebrum, which is our main interest here. Hence, specifically, for the CAP analysis, we used a mask excluding the cerebellum. We assessed temporal fluctuations of neural activity in terms of instantaneous BOLD CAPs following an approach by Gutierrez-Barragan et al. (2019) using a combined image-series from both WT and HET animals for each time point separately. Specifically, we first concatenated the processed image volumes from each animal from both genotypes to form a combined image-series. Then, we transformed this image-series into N, m-dimensional vectors where N is the total number of frames from all subjects and m is the total number of voxels within the mask mentioned above. We then divided all time frames using a K-means++ algorithm into clusters showing similar spatial activation patterns. We did this by assessing the spatial dissimilarity of the whole-brain activation pattern of each frame with that of every other frame using the correlation distance (1 - Pearson’s correlation) between every pair of m-dimensional vectors. The algorithm partitions the vector set into *K* clusters such that the sum of within cluster-distance 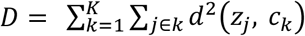 is minimized. Here, *K* is a predefined number of clusters and *d*(*z*_*j*_, *c*_*k*_) denotes the correlation distance between the centroid *c*_*k*_ of the *k*^*th*^ cluster and *j*^*th*^ time frame belonging to the *k*^*th*^ cluster. The k-means++ algorithm provides an optimal choice of initial cluster centroids as seeds so that distant centroids have a greater chance of getting selected as initial centroids. We varied the number of time frame clusters from 2 to 20, and in each case calculated the across-subject variance explained by the clustering solution as follows (Goutte et al., 1999):

- Within cluster variance, 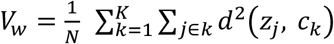 where, *N* is the total number of observations (time frames); *K* is number of clusters and *d*(*z*_*j*_, *c*_*k*_)denotes the correlation distance between the centroid *c*_*k*_ of the *k*^*th*^ cluster and *j*^*th*^ observation belonging to the *k*^*th*^ cluster.
- Between cluster variance, 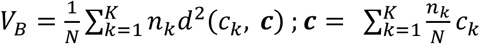 where *d*(*c*_*k*_, *c*) is the correlation distance between the global centroid ***c*** and the *k*^*th*^cluster’s centroid *c*_*k*_ and *n*_*k*_ is the number of time frames in the *k*^*th*^ cluster.
- Explained variance 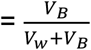

We then plotted the explained variance as a function of partitions of the image-series with increasing number of clusters (in the range from 2 to 20) and identified the minimum number at which the variance reached a saturation level (elbow point) as the optimal number of clusters. We also calculated, for each partition of k clusters, the fractional gain in explained variance over and above the variance explained by the partition of k-1 clusters. We further ensured the variance saturation at the elbow point by verifying that for all partitions with higher numbers of clusters this fractional gain in explained variance remained below 0.005 (Gutierrez-Barragan et al., 2019). Voxel-wise averaging of signal intensities was carried out across all time frames within each cluster to produce combined CAPs. Similarly, averaging across the genotype-specific time frames within each cluster was carried out to obtain the WT and the HET group-level CAPs.

We then performed a one-sample two-tailed T-test to test the mean activation, across the occurrences of each CAP in the combined image-series as well as the WT or HET sections of it, of each voxel against a null hypothesis of zero activation and corrected for multiple comparisons using the Bonferroni correction (p<0.01). The voxels that showed significant activation or deactivation constituted the one-sample T-statistic maps for each CAP, representing the regional activities that constitute the CAP. We then calculated the temporal and spatial properties for each of the CAPs for each subject within each genotype group:

1. Occurrence percentage: the percentage of the time frames corresponding to a CAP within a subject’s image-series.
2. Duration: number of consecutive frames corresponding to a CAP, averaged across all its occurrences, within a subject.
3. Voxel-level activations: Mean activation across all occurrences of a CAP within (a) either a genotype group-level image-series (WT/HET portions of combined image series) or (b) within a subject’s image-series.

### Regions of Interest (ROIs)

In order to compare the spatial properties of individual CAPs between the two genotypes, we obtained the mean activation in 28 bilateral ROIs in addition to voxel-level activation. These regions were selected based on their size (# of voxels > 30) and their relevance for the CAPs. These 28 anatomical ROIs were:

1. Default-mode like network (DMLN) regions: Cingulate (CgCtx), retrosplenial (RspCtx), and orbital (OrbCtx) cortices.
2. Associated cortical network (ACN) regions: Auditory (AuCtx) and Visual (VCtx) cortices.
3. Lateral cortical network (LCN) regions: Somatosensory (SSCtx), Motor (MCtx), Frontal Association (FrCtx) cortices and the Caudate putamen (CPu).
4. Other cortical and sub-cortical regions: Piriform (PirCtx), Rhinal (RhCtx), Thalamus (Th), Hippocampus (Hp) and the Olfactory bulb (OB).

### Statistical comparisons

We first compared, between genotypes at each time point, the median duration and occurrence percentage across subjects, for each of the optimal number of CAPs identified using the explained variance approach mentioned above. Mean activation levels of significantly activated (p < 0.01, 1-sample T-test, Bonferroni corrected) voxels in either the WT or the HET group for each of these CAPs across their occurrences were compared between genotypes using a 2-sample T-test and corrected for multiple comparisons using the Benjamini-Hochberg correction (Benjamini and Hochberg, 1995) for controlling the false discovery rate (FDR). The voxels that showed significant inter-genotype difference in activation constituted the 2-sample T-test statistic map and their T-statistics were displayed using two colormaps depending on whether they were activated or deactivated in the WT. We also compared mean, across subjects, activation for 28 ROIs between genotypes using a two-sample T-test and corrected using FDR. This ROI-level measure was calculated by taking a sum of one-sample T-test statistic values of all significantly activated (positive T) & deactivated (negative T) voxels belonging to the ROI and dividing it by the total number of voxels within that ROI, for every CAP in every subject.

In order to assess whether the inter-genotypic changes in temporal properties (duration and occurrence percentage) were robust, we considered 19 partitions of the combined image-series with progressively increasing number of CAPs ranging between 2 and 20. Then, for each partition with *K* CAPs, we compared, the median subject-level temporal properties in each of the *K* CAPs. Statistical comparisons were performed using the Wilcoxon two sample rank-sum test and were corrected within each partition using the Benjamini–Hochberg correction for controlling FDR.

## Classification

In order to classify HET animals from WT at each time point, we used a multinomial logistic regression (MLR) with regularization as a classifier. We trained the classifier using properties of CAPs on 80% of the subjects (n = 29) constituting the training set, chosen randomly but in a genotype-stratified manner, and tested its accuracy on the remaining 20% (n = 8) constituting the validation set. As all subjects were used to obtain group-level CAPs, information on validation set subjects could bias the classifier to predict them more accurately than otherwise possible. In order to avoid this bias, we modified the CAP extraction procedure to avoid cross-contamination during the classification procedure. For this, we first extracted *K* (*K = 2 to 20*) group-level CAPs (gCAP) from only the training set and then obtained the subject-level CAPs for each training and test-set subject as follows. First, we calculated spatial correlation between each of the *K* gCAPs and the image-series of every subject using only the significantly activated voxels in that gCAP. Then every time frame in subject’s image-series was assigned a label between 1 and *K* reflecting the gCAP with which it had the highest correlation. The frames with the same label were voxel-wise averaged to construct the corresponding CAP for every subject.

After obtaining *K* subject-level CAPs, we trained the classifier using their spatial and temporal properties in the training set subjects. Specifically, for each of the *K* CAPs we pooled either, (a) duration and occurrence rate (the temporal component), or (b) BOLD intensities of voxels whose activations within the corresponding gCAP were found to be significantly different from zero (the spatial component). These computed scores from each subject were z-scored within each subject so that their relative rankings, and not the absolute values, were used as input features for classification. The regularization parameter in the MLR classifier was set to 10 to control for over-fitting. This value was estimated using a grid search algorithm for estimation of hyper-parameters. The classifier was tested on the validation set and classification accuracy was obtained for every *K* between 2 and 20. We then shuffled the class identities (WT/HET) across all subjects and using the same training and test-set, obtained a chance level accuracy for every *K*. As we considered 19 partitions with the number of CAPs (*K*) ranging between 2 and 20, we obtained 19 real (actual class identities) and chance-level (shuffled class identities) accuracy values.

We repeated the accuracy calculation on 50 trials of randomly sampled, genotype stratified train and validation sets and statistically compared the median classification accuracy with median chance-level accuracy for all 19 values of *K*. As both the real and chance-level accuracy values for each split were obtained using identical training and test features (but different class identities), we performed this comparison using the Wilcoxon signed-rank test which allows comparison of paired samples. We corrected for 19 comparisons using FDR. Median chance-level accuracy was expected to be ∼ 50% as the number of subjects in each group was nearly equal.

## Results

### CAP identification

We started by identifying the optimal number of CAPs at each age. Figure 1 shows the total variance explained across subjects by each partition of the concatenated image-series from all subjects (18 WT and 19 HET). Visual identification of the elbow point of this curve that also ensured less than 0.5% gain in variance beyond it (bottom panels, Figure 1) showed partitions with 8, 3, and 6 clusters at the 3, 6, and 10-month time points, respectively, were optimal. As the three time points showed quite different explained variance versus number of clusters curves (in particular, the 6-month time point from the other two), we decided to perform the CAP analysis separately at each time point. CAPs were obtained by averaging the z-scored BOLD signals, voxel-wise, across all time frames with identical cluster membership.

**Figure 1:**
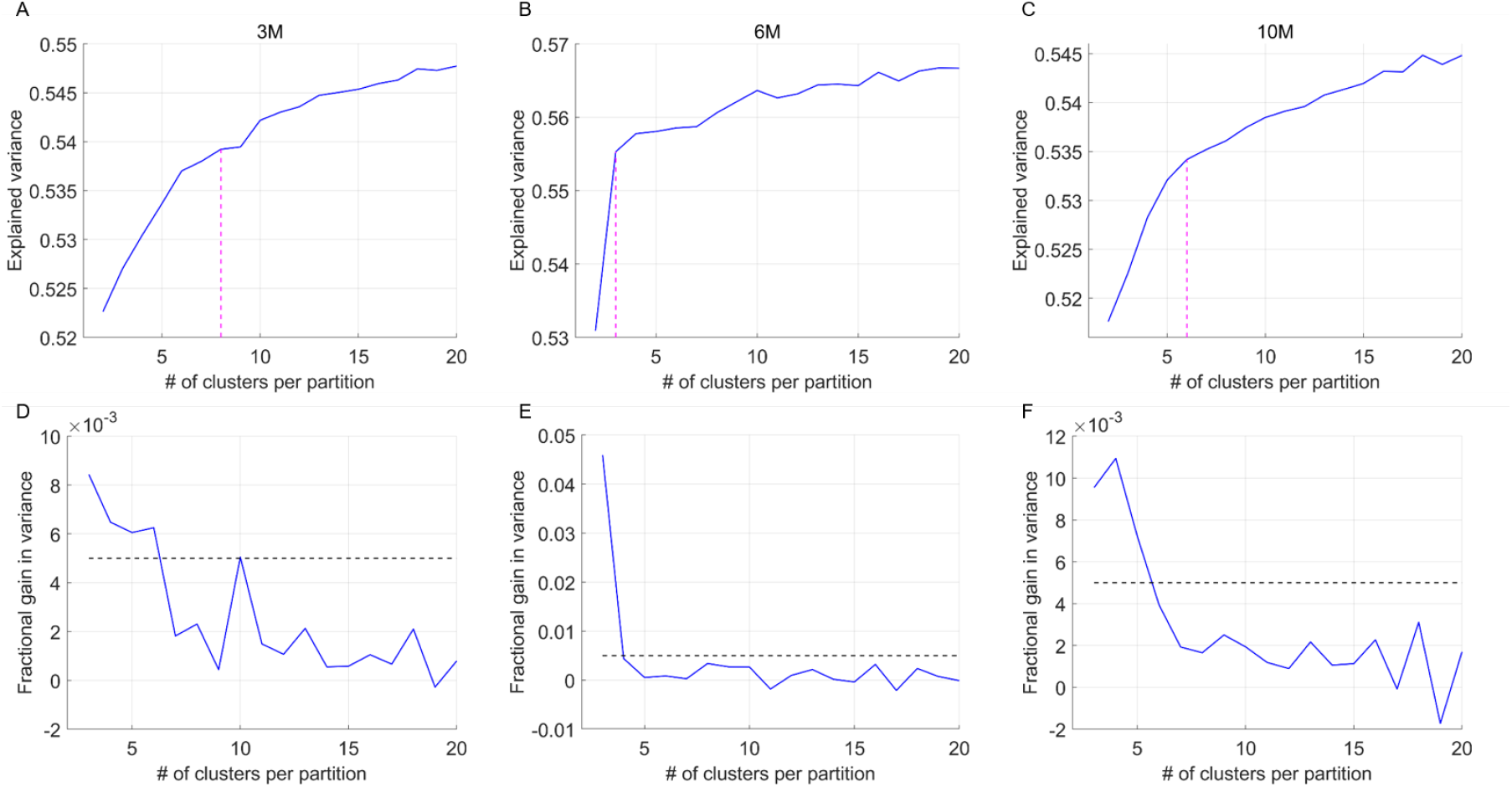
**A-C:** Across-subject variance explained as a function of number of clusters in the partition of the combined image-series at the 3-month (A), 6-month (B) and the 10-month (C) time points. Magenta dashed-line indicates the elbow point beyond which the explained variance is found to saturate in each case. We found 8, 3 and 6 clusters were sufficient to explain ∼53-55 % of variance at the 3, 6 and 10-month time points respectively. **D-F:** Fractional gain in the explained variance as the number of clusters in the partition increase from k to k+1, as a function of partitions, for 3-month (D), 6-month (E) and the 10-month (F) time points. We find that the fractional gain for every k after the elbow point is lower that 0.5%.

### Description of CAPs – temporal & spatial features

Figure 2 shows the first four of the 8 CAPs found at the 3-month time point. The genotype group-wise spatial pattern for each CAP (top panels, A, D, G, J) is represented by the color-coded t-value of voxels that show significant non-zero activation or deactivation (p < 0.01, one-sample T-test, Bonferroni corrected) within the WT or the HET group. The bottom panels of Figure 2A, 2D, 2G and 2J show the t-statistic values for voxels that show a significant inter-genotype difference (WT-HET) in activation (p < 0.05, 2-sample T-test, FDR corrected). The first CAP (LCN CAP; Figure 2A) is characterized by simultaneous co-deactivations of the mouse default-mode-like network (DMLN) composed of anterior cingulate, orbital, visual, retrosplenial, entorhinal cortices as well as dorsal hippocampus, and co-activations of lateral-cortical network (LCN) constituted by somatosensory, motor, and frontal association cortices as well as the caudate putamen. The spatial patterns of the second CAP (DMLN CAP, Figure 2D) showed a pattern anti-correlated with the LCN CAP. The remaining six CAPs could also be grouped in three spatially anti-correlated pattern pairs (Figures 2G, 2J, Supplementary Figure 1 A, D, G, J). CAPs 3 and 4 (Figures 2G and 2J) show widespread cortico-striatal co-deactivation and co-activation, respectively. Importantly, spatial topology of the first four CAPs is qualitatively very similar to those found by Gutierrez-Barragan et al., 2019. Figure 2B-C, 2E-F, 2H-I and 2K-L show the comparison of the distribution of subject-wise temporal properties – duration and occurrence percentage - of each CAP between the WT and HET group. Supplementary Figure 1 shows the spatial patterns and inter-group comparisons of spatial and temporal properties of four remaining CAPs identified at the 3-month time point. None of the CAPs showed any significant inter-genotype difference in either the duration or the occurrence percentage.

**Figure 2:**
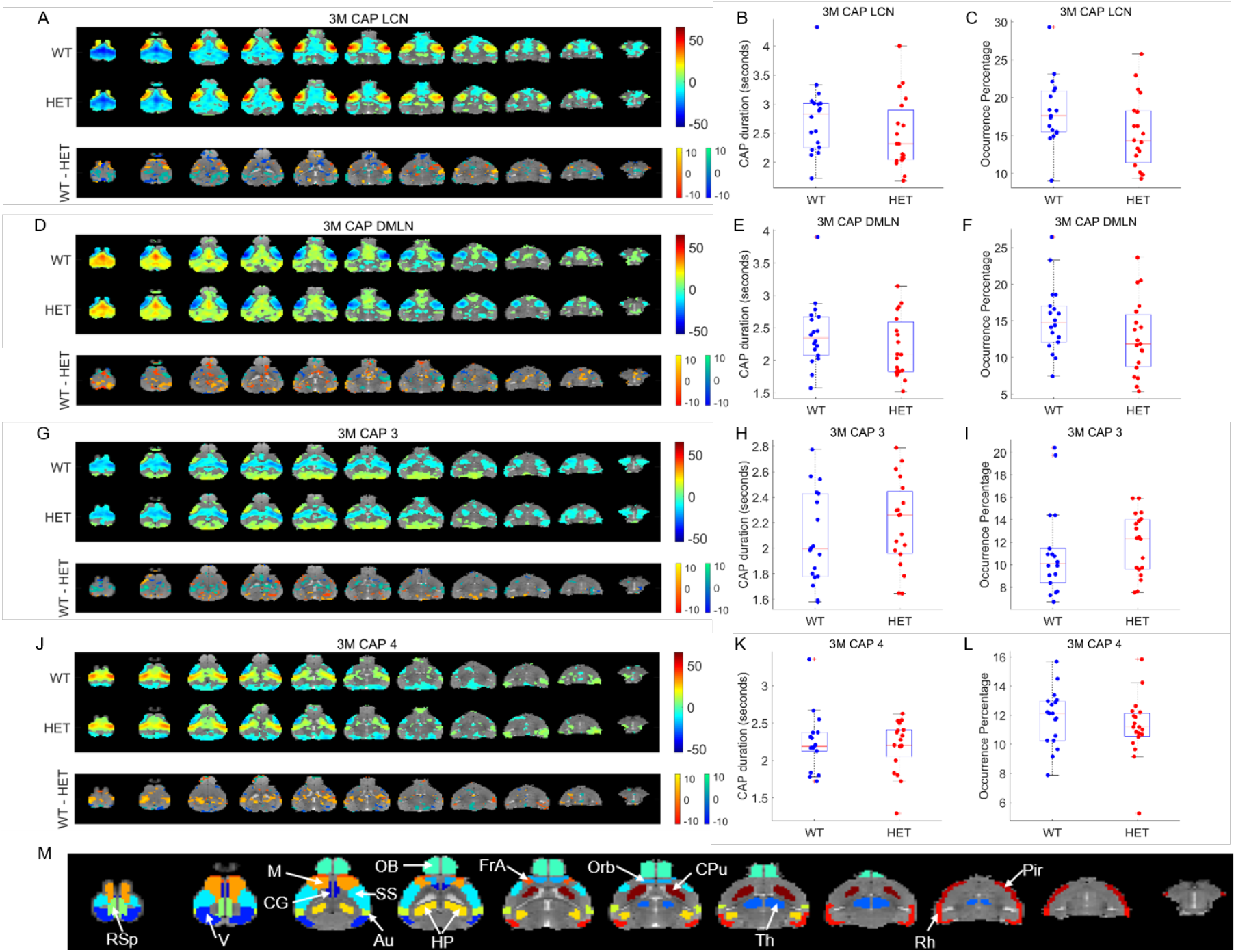
Comparison of spatial and temporal properties of 4 out of 8 CAPs at the 3-month time point. **A, D, G, J:** top panels show the one-sample T-statistic maps of significantly (Bonferroni corrected, p<0.01) activated and deactivated voxels for each CAP obtained from its occurrences in the WT and HET portions of the combined image-series. Bottom panels show the 2-sample T-test statistic map of voxels with significant (FDR corrected, p<0.05) difference in the (de)activation between the WT and HET CAPs. We make these comparisons for all voxels that are significantly activated or deactivated in either the WT or the HET group. Red-yellow and blue-green colour bars refer to voxels that are co-activated and co-deactivated respectively in the WT group. Thus, positive (yellow, green) and negative (red, blue) T-statistic values respectively indicate significantly lower and higher magnitude of activation in the HET group compared to the WT group. **B, E, H, and K:** Box-plots of comparisons of median, across subjects, duration of each CAP between WT & HET groups. **C, F, I, L:** Box-plots of comparison of median occurrence percentage, across subjects, of each CAP between WT & HET groups. **M:** Fourteen prominent ROIs considered in this study: Retrosplenial cortex (RSp), Visual cortex (V), Motor cortex (M), Cingulate cortex (CG), Olfactory bulb (OB), Somatosensory cortex (SS), Auditory cortex (V), Hippocampus (H), Frontal Association cortex (FrA), Orbital cortex (Orb), Caudate Putamen (CPu), Thalamus (Th), Rhinal cortex (Rh), and Piriform cortex (Pir).

Figure 3 shows the spatial patterns in the WT & HET group as well as inter-group comparison of spatial and temporal properties of the three CAPs identified at the 6-month time point. As in the case of the 3-month time point, the first two CAPs were characterized by simultaneous co-deactivation of DMLN and co-activation of LCN and vice versa, respectively. CAP 3 showed the widespread co-deactivation of cortical-striatal regions. None of the three CAPs showed any significant difference in either of the two temporal properties when corrected for 3 comparisons using FDR, however, the DMLN CAP showed a trend towards reduced median duration in the HET group in comparison with WT (p = 0.017, uncorrected). Differences in the magnitudes of activation and deactivation were found in several voxels for each of the three CAPs as evidenced from the 2-sample T-test maps (bottom panels, Figures 3A, D, G). These differences are quantified and detailed in the next section.

**Figure 3:**
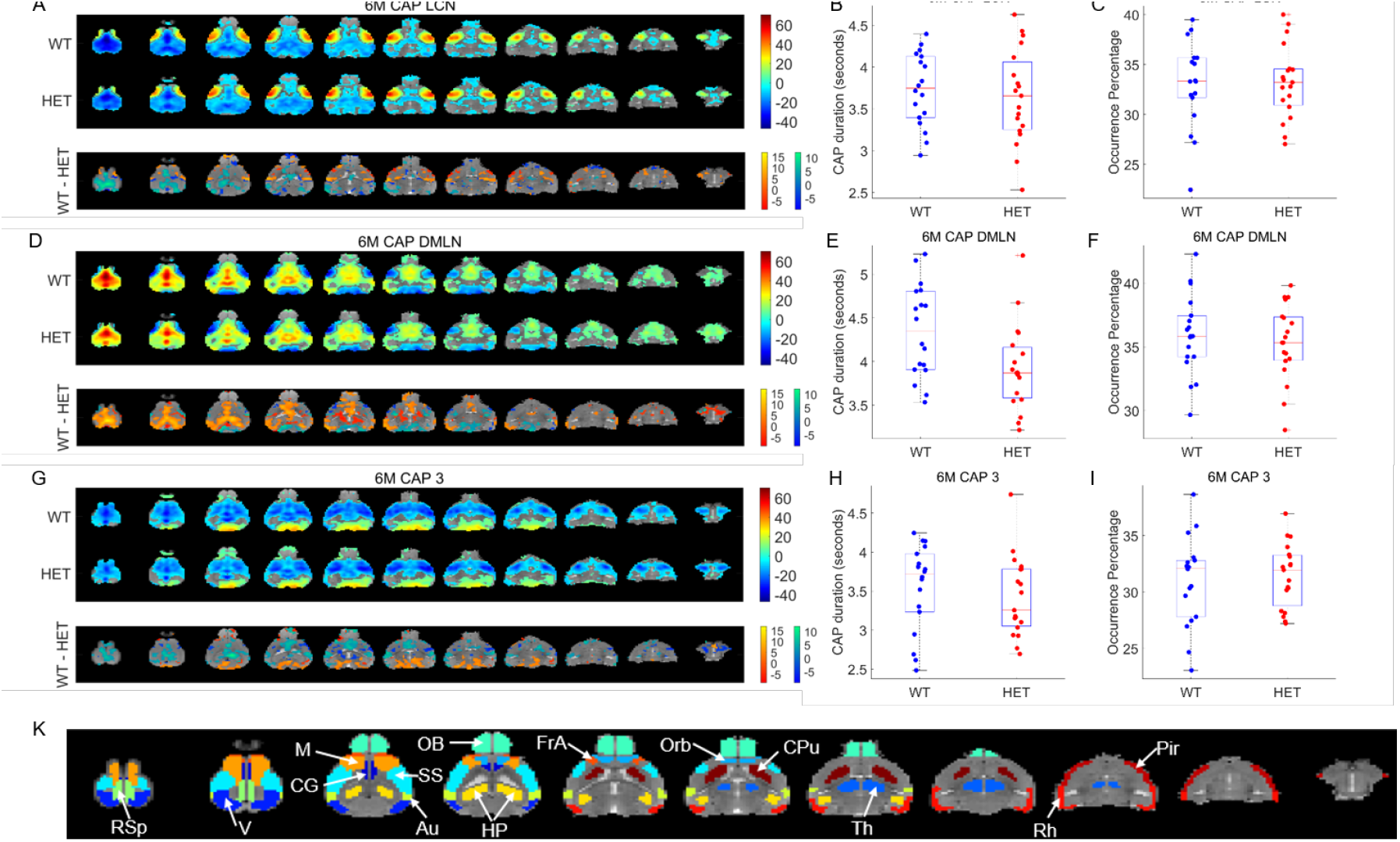
Comparison of spatial and temporal properties of 3 CAPs at the 6-month time point. **A, D, G:** top panels show the one-sample T-statistic maps of significantly (Bonferroni corrected, p<0.01) activated and deactivated voxels for each CAP obtained from its occurrences in the WT and HET portions of the combined image-series. Bottom panels show the 2-sample T-test statistic map of voxels with significant (FDR corrected, p<0.05) difference in the (de)activation between the WT and HET CAPs. We make these comparisons for all voxels that are significantly activated or deactivated in either the WT or the HET group. Red-yellow and blue-green colour bars refer to voxels that are co-activated and co-deactivated respectively in the WT group. Thus, positive (yellow, green) and negative (red, blue) T-statistic values respectively indicate significantly lower and higher magnitude of activation in the HET group compared to the WT group. **B, E, H:** Box-plots of comparisons of median duration, across subjects, of each CAP between WT & HET groups. **C, F, I:** Box-plots of comparison of median occurrence percentage, across subjects, of each CAP between WT & HET groups. **K:** Fourteen prominent ROIs considered in this study: Retrosplenial cortex (RSp), Visual cortex (V), Motor cortex (M), Cingulate cortex (CG), Olfactory bulb (OB), Somatosensory cortex (SS), Auditory cortex (V), Hippocampus (H), Frontal Association cortex (FrA), Orbital cortex (Orb), Caudate Putamen (CPu), Thalamus (Th), Rhinal cortex (Rh), and Piriform cortex (Pir).

At the 10-month time point, the LCN and the DMLN CAPs showed significant reduction in their median duration in the HET group (p < 0.05, Wilcoxon ranksum test, FDR corrected for 3 comparisons; Figures 4B, E). No other CAP showed any significant inter-genotype difference in the duration and the occurrence percentage. Two-sample T-test comparisons of spatial patterns between groups showed several prominent differences in the LCN and the DMLN CAPs (bottom panels, Figures 4A, 4D, 4M, 4P).

**Figure 4:**
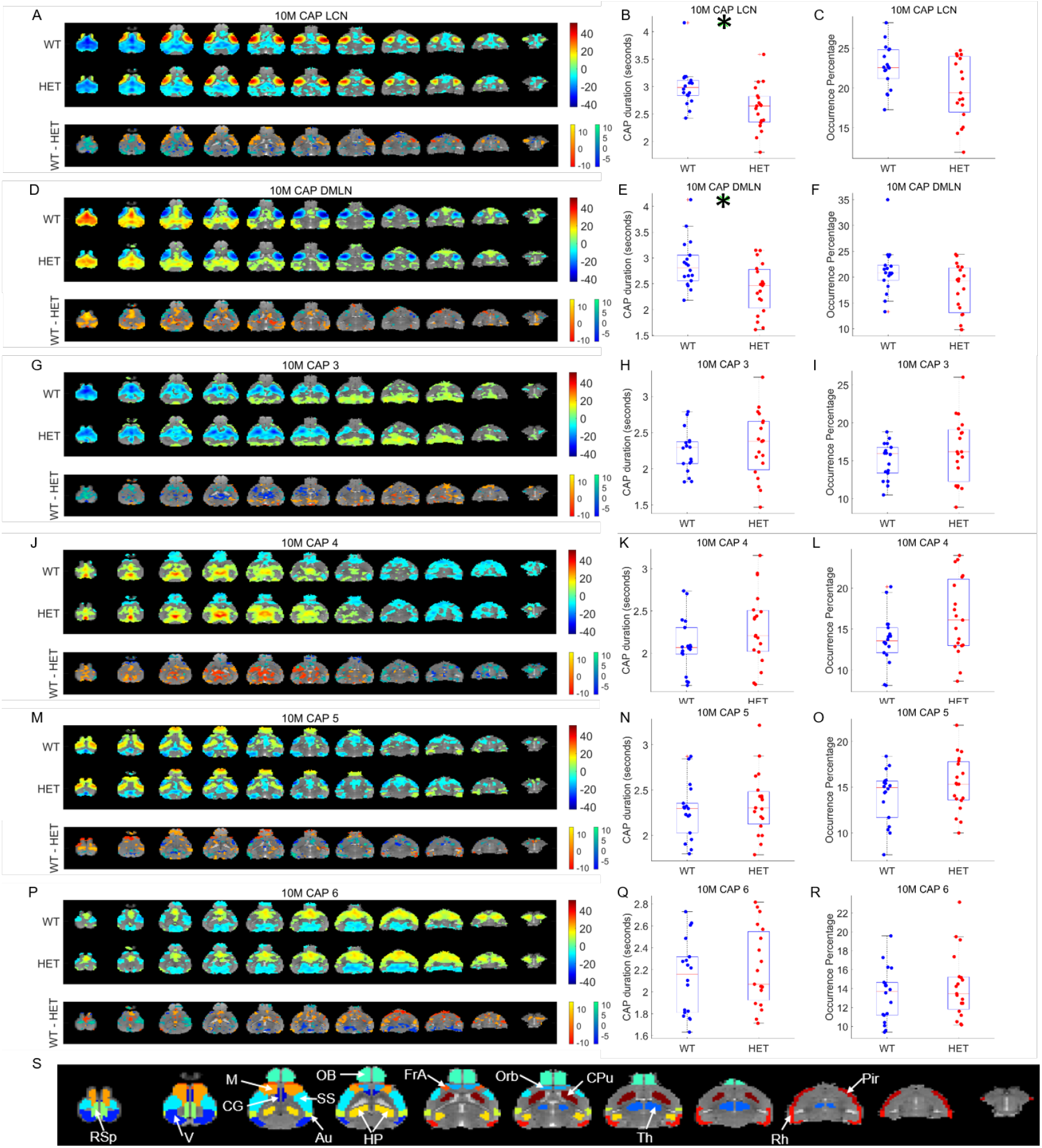
Comparison of spatial and temporal properties of 6 CAPs at the 10-month time point. **A, D, G, J, M, P:** top panels show the one-sample T-statistic maps of significantly (Bonferroni corrected, p<0.01) activated and deactivated voxels for each CAP obtained from its occurrences in the WT and HET portions of the combined image-series. Bottom panel shows the 2-sample T-test statistic map of voxels with significant (FDR corrected, p<0.05) difference in the (de)activation between the WT and HET CAPs. We make these comparisons for all voxels that are significantly activated or deactivated in either the WT or the HET group. Red-yellow and blue-green colour bars refer to voxels that are co-activated and co-deactivated respectively in the WT group. Thus, positive (yellow, green) and negative (red, blue) T-statistic values respectively indicate significantly lower and higher magnitude of activation in the HET group compared to the WT group. **B, E, H, K, N, Q:** Box-plots of comparisons of median duration, across subjects, of each CAP between WT & HET groups. **C, F, I, L, O, R:** Box-plots of comparison of median occurrence percentage, across subjects, of each CAP between WT & HET groups. Green asterix indicates significant inter-group difference (p < 0.05, Wilcoxon rank-sum test, FDR corrected for 6 comparisons). **S:** Fourteen prominent ROIs considered in this study: Retrosplenial cortex (RSp), Visual cortex (V), Motor cortex (M), Cingulate cortex (CG), Olfactory bulb (OB), Somatosensory cortex (SS), Auditory cortex (V), Hippocampus (H), Frontal Association cortex (FrA), Orbital cortex (Orb), Caudate Putamen (CPu), Thalamus (Th), Rhinal cortex (Rh), and Piriform cortex (Pir).

Although the optimal number of CAPs was different at each age, there was a high degree of similarity in the spatial patterns of activation and deactivation across ages, especially for the LCN and the DMLN CAPs (Supplementary Figure 2). The lowest correlation for the LCN CAP was 0.93 between the 3-month and the other two time points. The lowest correlation for the DMLN CAP was 0.68 between the 6- and the 10-month time points. The LCN-DMLN CAP anti-correlation was also observed at all time points with the lowest value of -0.68 observed between the 6-month DMLN and the 10-month LCN CAPs.

### Quantification of inter-genotype spatial differences between CAPs

Next, we quantified the extent of inter-genotype difference in the magnitudes of co-activation and co-deactivation in terms of percentages of all voxels from 28 ROIs. We divided the voxels in each region in three categories for each CAP: (i) co-activated voxels in WT, (ii) co-deactivated voxels in WT and (iii) non-activated voxels in either group. Categories (i) and (ii) were each subdivided into 3 sub-categories that showed significant increase, decrease, and non-significant difference in the activation and deactivation magnitude, respectively, in the WT group in comparison with the HET group.

Figure 5 shows the distributions of all voxels in these 28 brain regions according to the seven categories mentioned above for the LCN and the DMLN CAP at the three time points. Supplementary Figure 3 shows similar distributions for other CAPs. At all three time points, DMLN regions (cingulate and retrosplenial cortices, in particular) are co-activated during the DMLN CAP and co-deactivated during the LCN CAP. The LCN regions (somatosensory and motor cortices, in particular), on the other hand, include both significantly (p < 0.01; Bonferroni corrected) activated as well as deactivated voxels during both CAPs. In the 3-month LCN CAP, majority of activated voxels showed lower deactivation in the right visual and auditory cortices while increased deactivation in the right orbital cortex in the HET group. Prominent changes in the DMLN CAP include lower and higher activation magnitudes in HET in the retrosplenial and cingulate cortices respectively in both hemispheres (Figure 5A, 5B). At the 6-month time point, all DMLN regions consistently showed reduced magnitudes of activation and deactivation in HET in the DMLN and LCN CAPs respectively (Figure 5C, 5D). Most voxels from the LCN regions also showed reduced magnitude of activation and deactivation in both CAPs in the HET group. Similar reductions in activation magnitudes were also found in a majority of voxels from the LCN and the DMLN regions in both CAPs in the HET group at the 10-month time point (Figure 5E, 5F).

**Figure 5:**
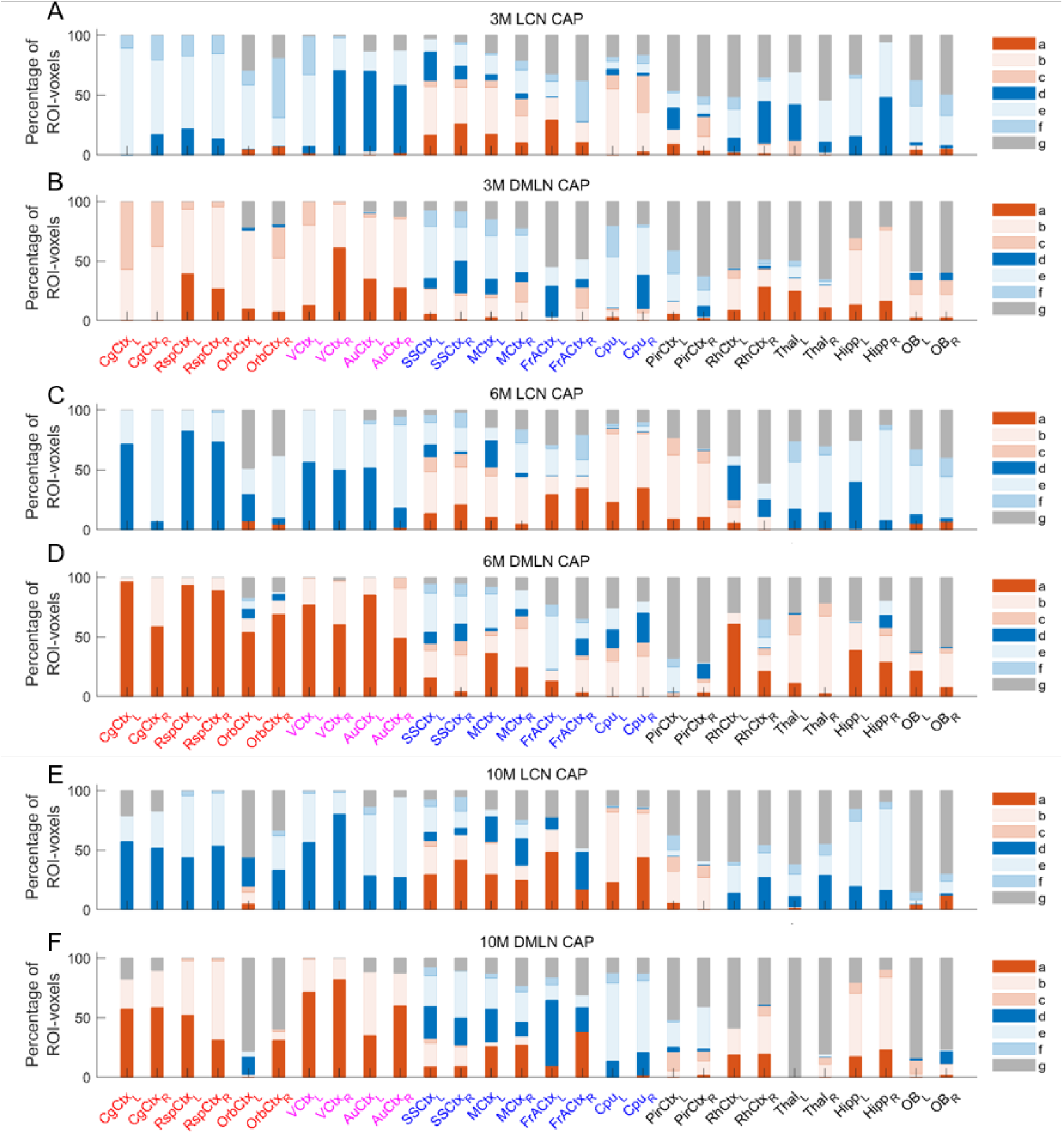
Percentages of voxels per ROI belonging to one of 7 categories for the LCN CAP (A (3M), C(6M), E(10M)) and the DMLN CAP (B (3M), D(6M), F(10M)) respectively. The red & blue colour bars indicate significant co-activation and co-deactivation, respectively, in either the WT or the HET group. Different colour/shade bars indicate percentage of voxels from seven different categories: (a) significant co-activation and higher activation magnitude in WT, (b) significant co-activation and no significant inter-genotype difference in the activation magnitude, (c) significant co-activation and higher activation magnitude in HET, (d) significant co-deactivation and higher activation magnitude in WT, (e) significant co-deactivation and no significant inter-genotype difference in the activation magnitude, (f) significant co-deactivation and higher activation magnitude in HET, and (g) non-significant co-activation or co-deactivation during a CAP. Voxel-wise one-sample T-test (p < 0.01, Bonferroni corrected) and two-sample T-test (p < 0.05, FDR corrected) are performed across occurrences of a CAP within the concatenated genotypic image-series from all subjects to identify the voxels belonging to one of the 7 categories mentioned above in the entire brain. We then calculate the percentages of each category in each of the 28 ROIs shown in Figures 2M, 3K, 4S. ROIs belonging to the DMLN have red labels, ROIs belonging to the associated cortical network have magenta labels while the blue labels represent LCN ROIs. The black labels represent the rest of the cortical and sub-cortical ROIs.

While Figure 5 outlines proportions of co-activated and co-deactivated voxels in each ROI and what percentage of them show significant inter-genotype difference in magnitudes of co-(de)activation, it does not compare the actual activation levels between genotypes per region. Therefore, we calculated regional activation measure from the 1-sample T-test statistic map for each subject and compared the mean T-statistic of significant (de)activations per region, averaged across subjects, between the genotypes. Figure 6 shows an inter-genotype comparison of mean +/-SEM of the weighted mean of T-statistic per region, averaged across subjects, for LCN and DMLN CAPs. We found no significant differences in the mean T-statistics at the 3-month time point for either the LCN or the DMLN CAP (Figure 6A, 6B). Significant reductions were found for some regions in the DMLN CAP at both the 6- and the 10-month time points (Black asterix, Figure 6C, 6D, 6E, 6F).

**Figure 6:**
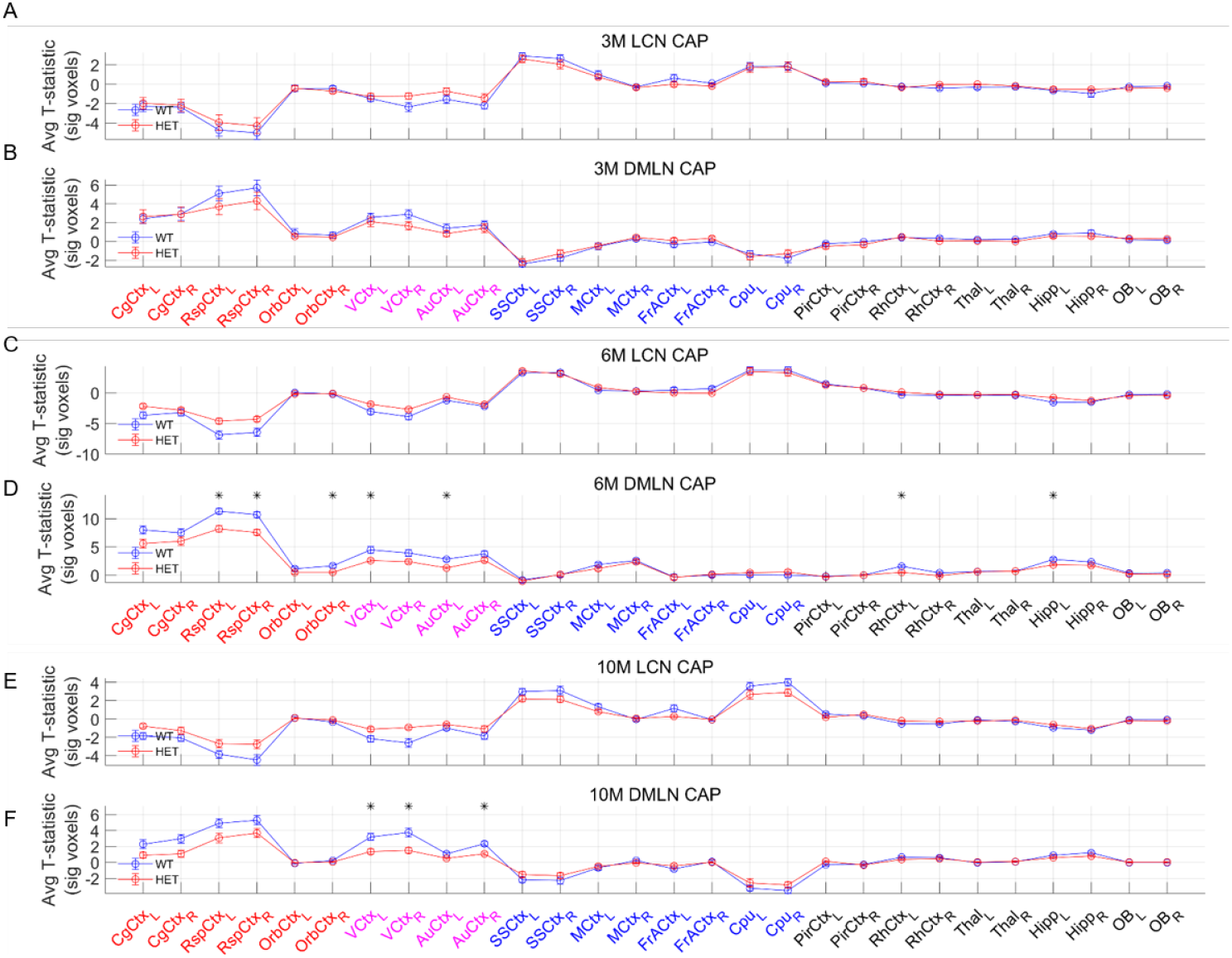
Comparison of mean activation per ROI across subjects for the LCN (panels A, C, E) and the DMLN CAP (B, D, F). Mean activation per ROI at the subject level is calculated by first identifying significantly co-activated or co-deactivated voxels using a one-sample T-test (p < 0.05, Bonferroni corrected) performed across occurrences of a CAP within the subject’s scan and then dividing the total T-test statistic from those voxels by the total number of voxels within each ROI. Across-genotype comparisons were performed using a two-sample T-test corrected for multiple comparisons using false discovery rate. Black asterix shows significant differences between genotypes (p < 0.05, FDR corrected for 28 comparisons). ROIs belonging to the DMLN have red labels, ROIs belonging to the associated cortical network have magenta labels while the blue labels represent LCN ROIs. The black labels represent the rest of the cortical and sub-cortical ROIs.

### Robustness of temporal alterations in CAPs

As mentioned before, we identified CAPs after finding an optimal partition of the combined image-series that explained a saturation-level variance across subjects. While a clear elbow point was visible at the 6-month time point (Figure 1B), the explained-variance increased monotonically with the # of clusters at the 3- and 10-month time points. Therefore, we aimed to assess whether the inter-genotype change in the temporal properties found in some CAPs, especially at the 10-month time point, could be observed consistently when the image-series is partitioned in different numbers of clusters. Hence, we compared the median duration and occurrence percentage for each of the *K* CAPs with *K* ranging between 2 and 20. Figure 7 shows the p-values of comparison of median durations (Figure 7A-C) and the occurrence percentage (Figure 7D-F) for each CAP (black marker) as a function of *K*. Since the null-hypotheses tested across values of *K* are not necessarily independent (spatially similar CAPs can be identified across different *K*), the p-values should be corrected for the number of comparisons within each partition. Orange circles indicate the critical p-values for Bonferroni correction within the partition and hence CAPs with p-value close to or lower than these critical values can be considered to show significant inter-genotype difference. Only a few CAPs showed significant inter-genotype difference in either of the two temporal properties at the 3- and 6-month time points (Figure 7A-B, 7D-E). In contrast, several CAPs across multiple partitions showed significant inter-group differences in duration and/or occurrence at the 10-month time point (Figure 7C, 7F). Thus, the alterations in temporal properties of CAPs were consistent and robust across the choice of *K* at the 10-month time point.

**Figure 7:**
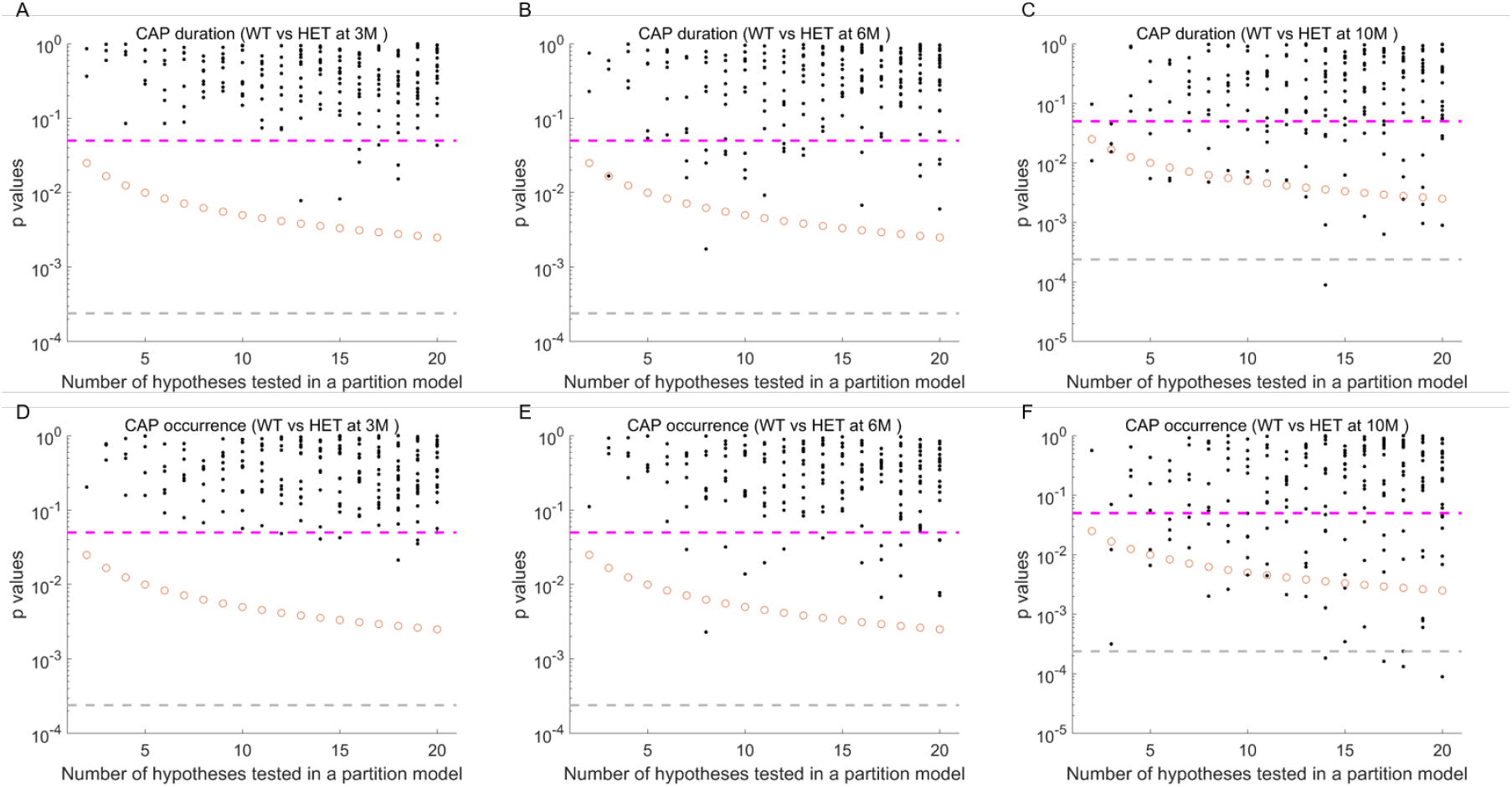
P-values of comparison of median duration (top panels) and occurrence fractions (bottom panels) between WT and HET for each CAP (black markers) from 19 partitions, with the number of clusters ranging from 2 to 20, of the combined image series with RS-fMRI data from both groups. Magenta dashed line: p=0.05, uncorrected; orange circle: Bonferroni corrected threshold p-value for each partition; grey dashed line: Bonferroni corrected threshold p-value across all partitions.

### Classification

Finally, we assessed the predictive power of CAPs in distinguishing the HET animals from the WT. Classification based on the temporal properties of CAPs did not perform better than chance at any of the three time points. Classification based on spatial features, however, performed with significantly higher accuracy than chance at all the three time points for most values of *K* (Figure 8; p < 0.05, unpaired T-test, FDR corrected for 19 comparisons). The accuracy at the 3-month time point was around 60% and increased to 80% at the 6- and 10-month time points. These accuracy levels reflect the robustness of differences in magnitudes of voxel-level activation between the two genotypes especially at the early and the late manifest states in the progression of the disease.

**Figure 8:**
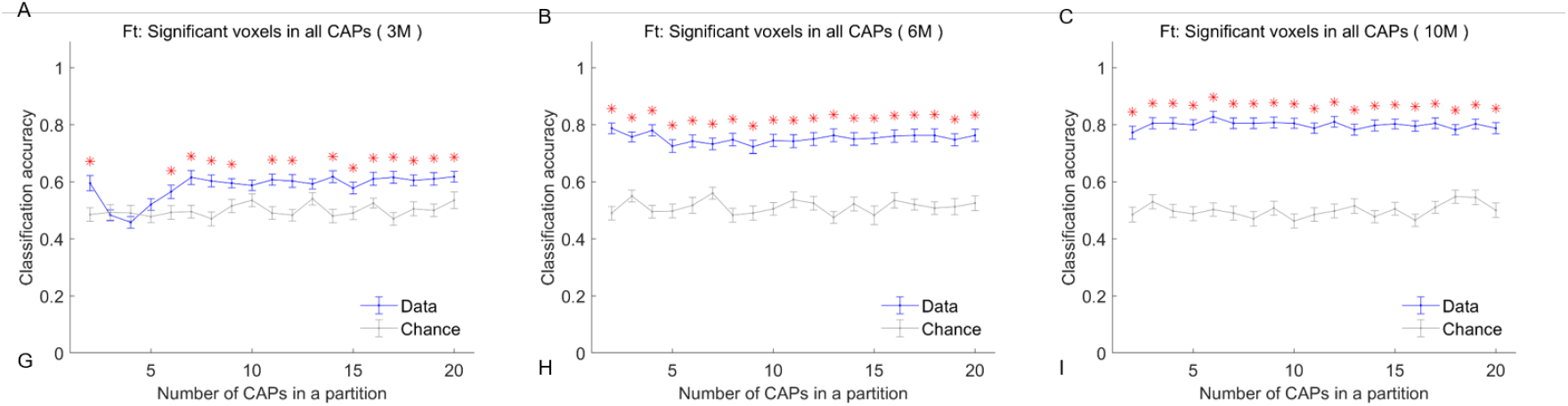
Classification accuracy (blue, mean +/-SEM) using z-scored BOLD signals of voxels with significant activations in all CAPs within a partition as a function of partitions of the combined image series at the 3-month (A), 6-month (B) & the 10-month (C) time point. The grey curve shows the corresponding chance-level accuracy (mean +/-SEM) & red asterisk indicates significantly higher mean accuracy than the chance level, after correcting for all 19 comparisons (p < 0.05; FDR corrected) in each panel.

## Discussion

In this longitudinal study, we investigated changes in the transient brain states constituting the resting-state of the brain in a mouse model of HD. We used the zQ175 knock-in DN HET model that shows molecular, cellular, and circuitry alterations that resemble those observed in PwHD (Heikkinen et al., 2020; Menalled et al., 2012; Southwell et al., 2016). The transient brain states were uncovered using the methodology of co-activation patterns that are constituents of resting-state networks. We found that the duration of two prominent, spatially anti-correlated CAPs was reduced in the HET group at the late manifest progression state. The first CAP, referred to as the lateral-cortical network (LCN) CAP, was characterized by simultaneous co-activation and co-deactivation of brain regions constituting the LCN and the default mode like network (DMLN), respectively. The second CAP, called the DMLN CAP, was characterized by the opposite activation topology. At the early-manifest state, the DMLN CAP showed a trend for reduction in average duration but the difference was not significant upon correcting for multiple comparisons. None of the CAPs at the premanifest state showed any significant difference in their temporal properties. In terms of spatial properties, magnitude of activation and deactivation was reduced in a large proportion of voxels of the prominent regions belonging to the LCN (somatosensory and motor cortices) and DMLN (anterior cingulate, retrosplenial, visual cortices) CAPs in the HET group at the early and late manifest states. At the premanifest state, a sizeable proportion of voxels in the orbital and the cingulate cortices showed higher activation/deactivation magnitude in the HET group in both LCN as well as DMLN CAPs. Since we could find statistically significant differences especially in the spatial properties of CAPs at all three states, we investigated their predictive power to distinguish HET animals from the WT. Here, we found that the spatial component of CAPs consisting of BOLD signals of significantly (de-)activated voxels in each CAP was 60% accurate in correctly predicting the genotypic identity of each individual test-set subject in each of the 50 trials at the premanifest state. At the early-, and late manifest states, the accuracy increased to 80% reflecting more widespread, robust, and regional level changes occurring as a result of phenotypic progression.

### Methodological considerations

We obtained CAPs using a recently improved method (Gutierrez-Barragan et al., 2019) that allows clustering of all frames instead of previous approaches where a fraction (typically 15%) of all frames corresponding to supra-threshold BOLD signal of a seed region were clustered (Karahanoğlu and Ville, 2015; Liu and Duyn, 2013a). This improved method allows a better estimate of temporal properties of CAPs since all frames are partitioned. Here, we concatenated the images of individual mice from both the WT and the HET group in a single image-series before partitioning it to identify the combined CAPs. The advantage of this approach over the separate group-level analysis lies in a joint search for common CAPs in all animals, enabling a direct comparison between groups without a need to match CAPs between groups. In our study, we regressed out the global signal before carrying out the CAP analysis. A previous CAP study in humans has reported results both with and without global signal regression (GSR) and they showed that GSR had little effect on the DMN CAPs involving simultaneous co-activation of DMN regions and co-deactivation of task-positive network (similar to LCN in mice) regions (Liu and Duyn, 2013b). Since the major differences we found in this study involved an anti-correlated pair of DMLN and LCN CAPs, GSR, per se, is unlikely to be a confound.

### Spatial topology of CAPs

At least four CAPs identified at the premanifest and late manifest states and all three CAPs at the early manifest states were similar to those previously identified (Gutierrez-Barragan et al., 2019). They also reported that the CAPs occurred in pairs of anti-correlated spatial patterns, a finding that was replicated here. The first pair demonstrated simultaneous co-activation of LCN regions such as somatosensory and motor cortices and co-deactivation of DMLN regions that included primarily anterior cingulate, retrosplenial, visual and orbital cortices, and vice versa. Another pair demonstrated widespread cortico-striatal activations and deactivations (only the deactivated pattern was isolated as a CAP at the 6-month time point as the image-series was partitioned into 3 clusters). Pearson’s correlation between spatial patterns of CAPs (z-scored BOLD signals) identified at the different ages was high, especially in the case of LCN and DMLN patterns.

### CAP changes in HD

In terms of temporal properties of duration and occurrence rates, the duration of the DMLN CAP showed a trend towards a decrease in the HET group at the early manifest state. Both the LCN and DMLN CAPs showed significantly reduced duration in the HET group at the late manifest state. When we compared the temporal properties of CAPs identified in different partitions of the image-series with the number of clusters ranging between 2 and 20, only a few CAPs across partitions showed significant inter-genotype differences at the pre- and early manifest states. At the late manifest state, though, a much higher number of CAPs showed significant alterations between genotypes. Thus, the changes in the temporal component of CAPs were most robust at the late manifest state compared to the earlier states considered in this study.

Spatially, we compared the activation strengths of different regions within each CAP between the two groups. The magnitude of activation/deactivation of a large percentage of voxels in the cingulate and the retrosplenial cortices, prominent regions in the DMLN, was found to be higher in the HET group at the premanifest state. At the early and late manifest states, however, we found a significant reduction in the magnitude of activation or deactivation in majority of voxels in prominent regions belonging especially to the DMLN and the LCN, in the HET group. A recent article shows that the activations in DMLN CAP in mice is correlated with a prominent unimodal-transmodal cortical structural connectivity gradient (Coletta et al., 2020). Therefore, while we cannot rule out vasculature changes in these regions as an underlying factor for the reductions in the magnitude of activation of both the DMLN and LCN CAPs, they could also be explained by potential changes in the underlying structural connectivity gradients in this mouse model

### Potential of RS-CAPs as early biomarkers of HD

Resting-state fMRI has been used to identify robust markers of several neurological and neurodegenerative diseases (Adhikari et al., 2017; Badhwar et al., 2017; Di Perri et al., 2018; Siegel et al., 2016). One of the prime reasons for this is the non-invasive nature of the scanning protocol. More importantly, task positive networks (TPNs) that are active in specific behavioral tasks such as the motor, attention, language networks are also active during the resting-state (Vincent et al., 2007). Hence, alterations within and inter-RSN functional connectivity can be used and has been useful in predicting behavioral deficits in humans (Ramsey et al., 2016; Siegel et al., 2016; Tavor et al., 2016). Changes in the static FC of DMN have been observed in AD patients (Mevel et al., 2011) and found to correlate with levels of amyloid-beta plaque deposits in regions of DMN (Koch et al., 2015; Myers et al., 2014; Sperling et al., 2009). RSNs such as the default mode and task-positive networks have rodent analogs (Gozzi and Schwarz, 2016), which means preclinical resting-state fMRI studies in rodent models of disease have high translational value. Going beyond the static FC, Belloy et al. investigated its temporal fluctuations in a transgenic mouse model of AD. They found that short (3s), spatio-temporal patterns of recurring neural activity, called the quasi-periodic patterns (QPPs), were altered in transgenic mice and that they contributed to the observed static FC changes (Belloy et al., 2018b). In the same cohort it was shown that RS-CAPs can predict the identity of transgenic animals accurately (Adhikari et al., 2021).

Development of monitoring and response biomarkers is a pressing need in HD research. Most clinical investigations on the impact of HD on the resting-state have focused on the static functional architecture (Pini et al., 2020). Preclinical RS-fMRI studies in mouse models of HD have also investigated changes only in static FC (Li et al., 2017). To our knowledge, our study is the first investigation of changes in the transient brain states that make up the resting brain in a mouse model of HD. We find that physiologically prominent brain-states constituting two most robustly observed RSNs – the default-mode like network and the lateral cortical network, and forming the major functional connectivity gradients show the most significant alterations in this mouse model and the changes can be observed from the premanifest state. The spatiotemporal changes in RS-CAPs at all three HD states considered in this study prompted us to investigate their predictive power in identifying the status of the HD mouse model. Here, we found nearly 80% accuracy in distinguishing the WT animals from the heterozygous animals when spatial features of CAPs were used at the early and late manifest states. The accuracy was expectedly lower at the premanifest state, although the classifier performance was significantly better than the chance level.

Our findings therefore suggest that we can consider RS-CAPs as sensitive markers in this HD mouse model starting from a premanifest state. This is crucial since studies involving animal models of the disease are carried out to identify early markers which is seldom possible in humans. RSNs include spatially distributed brain regions that show high functional connectivity obtained from the entire scan and analogous RSNs have been found in different species. RS-CAPs measure dynamic co-activations and co-deactivations of regions occurring during the scan, thus at shorter time scales than the RSNs. However, similar to RSNs, RS-CAPs - especially a pair of anti-correlated CAPs involving simultaneous co-activations and co-deactivations of default-mode like and lateral-cortical networks - have been identified across species including humans (Liu et al., 2018). Therefore, investigation of changes in properties of CAPs in a disease context has potential translational value. Further, as there is high degree of similarity in functional networks active during behavioral tasks and the resting-state, we can speculate that changes in RS-CAPs may be predictive of functional changes at the behavioral level. Therefore, findings from our study make a strong case to investigate RS-CAPs in a clinical setting in order to find out if they can be, first, predictive of the human disease and, next, predictive of behavioral impairments and/or stages of the disease in patients.

## Figures

**Supplementary Figure 1:**
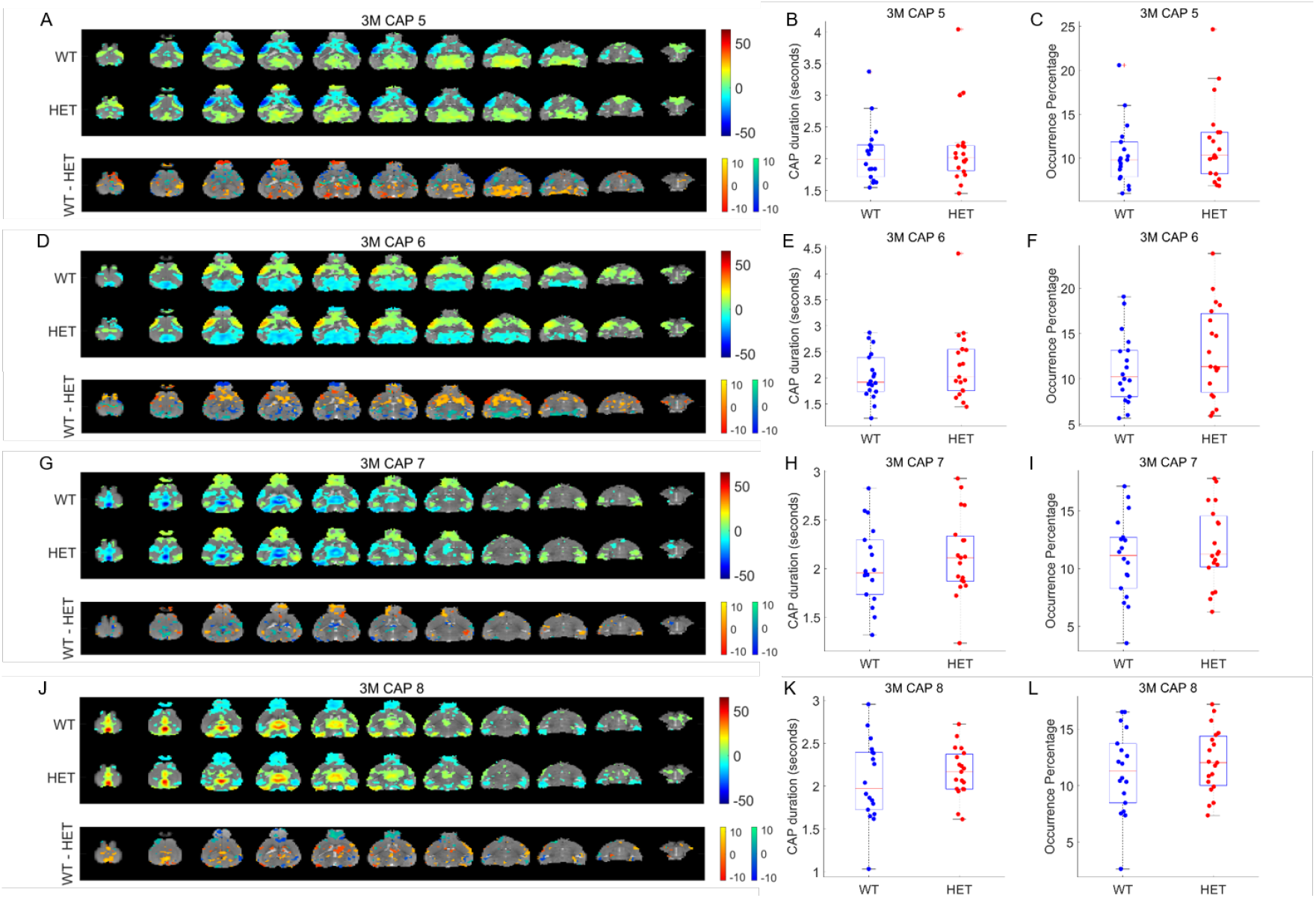
Comparison of spatial and temporal properties of the other 4 out of 8 CAPs at the 3-month time point. **A, D, G, J:** top panels show the one-sample T-statistic maps of significantly (Bonferroni corrected, p<0.01) activated and deactivated voxels for each CAP obtained from its occurrences in the WT and HET portions of the combined image-series. Bottom panels show the 2-sample T-test statistic map of voxels with significant (FDR corrected, p<0.05) difference in the (de)activation between the WT and HET CAPs. We make these comparisons for all voxels that are significantly activated or deactivated in either the WT or the HET group. Red-yellow and blue-green colour bars refer to voxels that are co-activated and co-deactivated respectively in the WT group. Thus, positive (yellow, green) and negative (red, blue) T-statistic values respectively indicate significantly lower and higher magnitude of activation in the HET group compared to the WT group. **B, E, H, K:** Box-plots of comparisons of median duration, across subjects, of each CAP between WT & HET groups. **C, F, I, L:** Box-plots of comparison of median occurrence percentage, across subjects, of each CAP between WT & HET groups. Black asterix indicates significant inter-group difference (p < 0.05, Wilcoxon rank-sum test, FDR corrected for 8 comparisons).

**Supplementary Figure 2:**
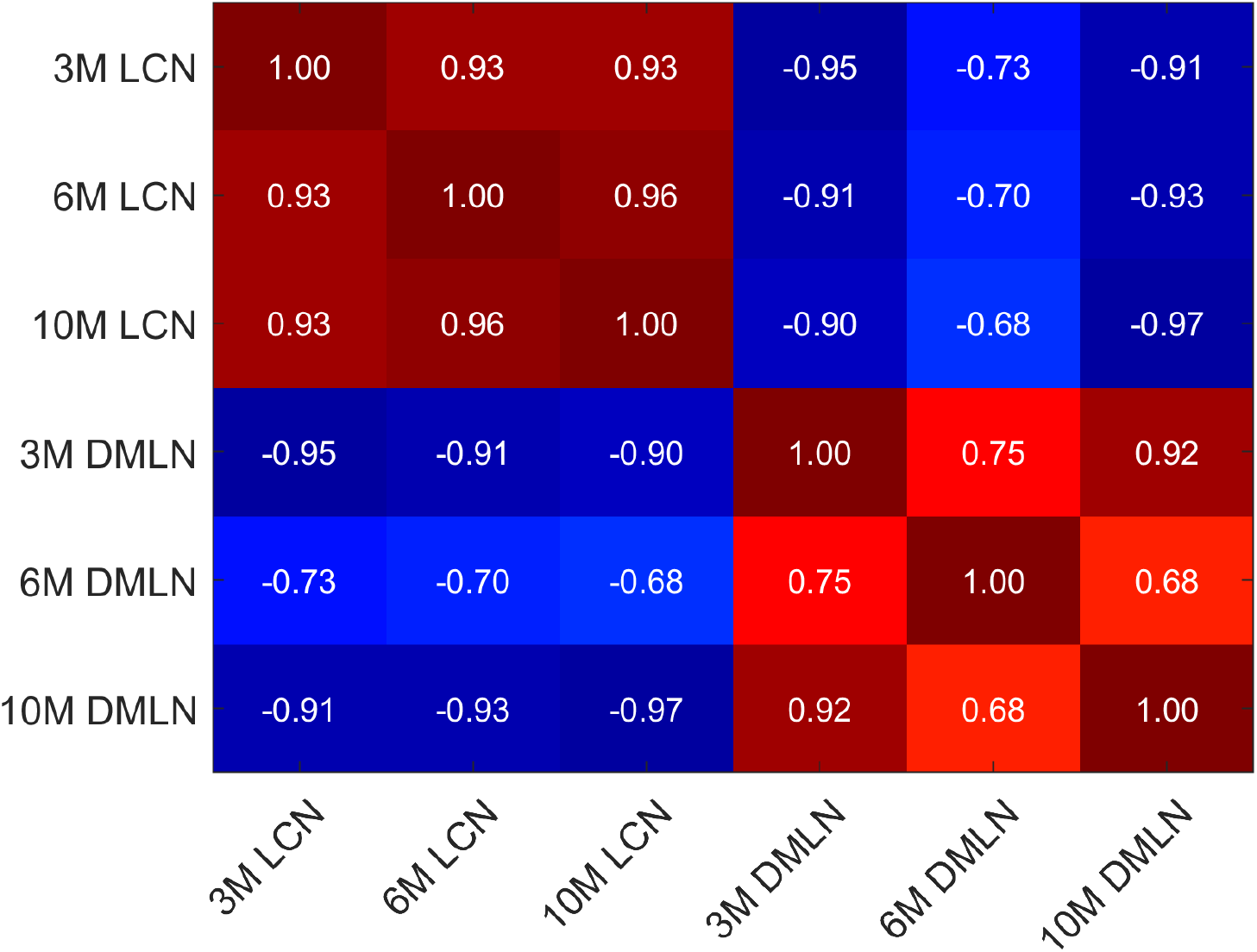
Pearson’s correlation coefficient between spatial patterns of the LCN and DMLN CAPs identified at three time points.

**Supplementary Figure 3:**
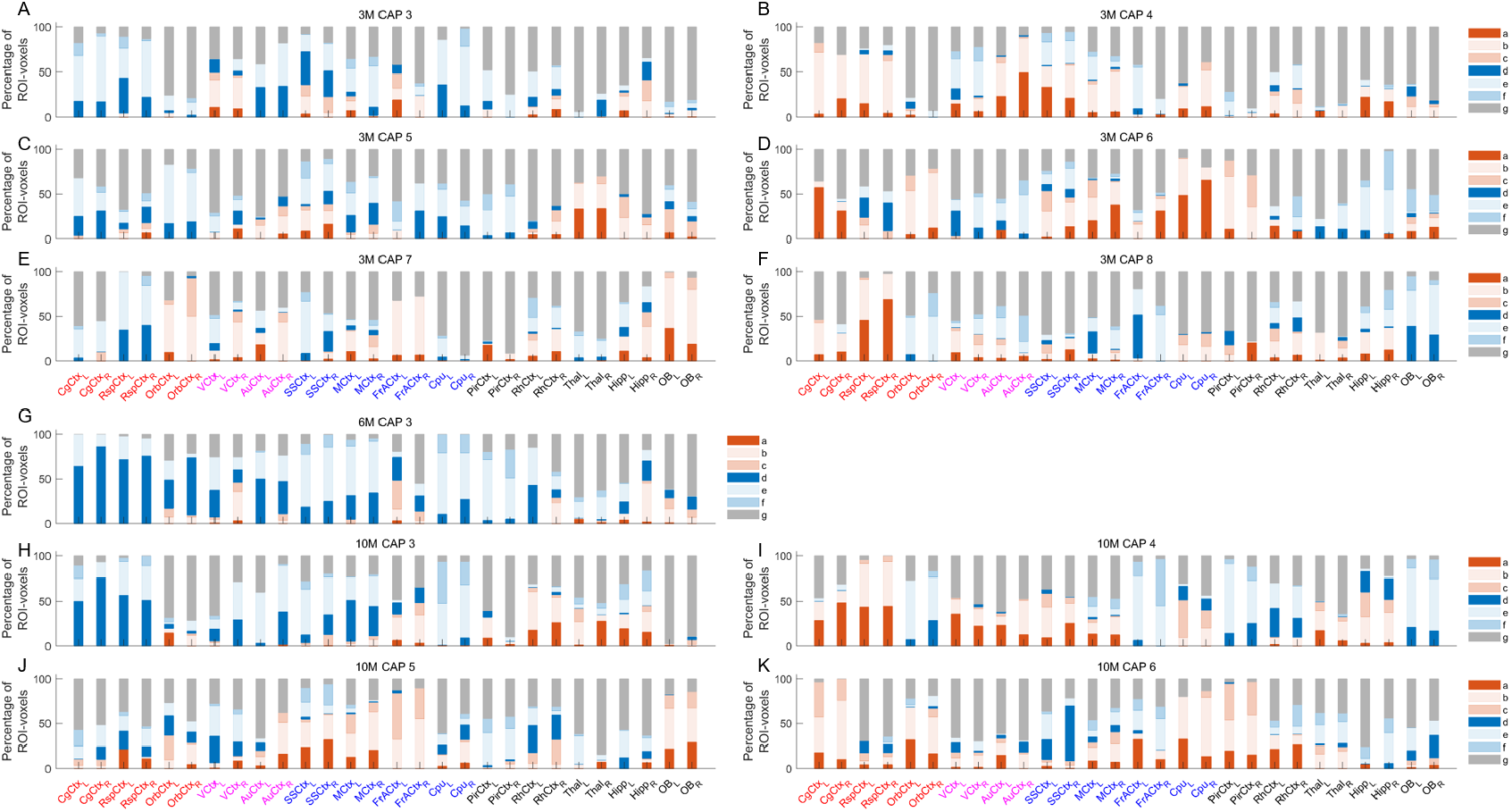
Percentages of voxels per ROI belonging to one of 7 categories for the other-than-LCN-DMLN CAPs (A-F (3M), G (6M), H-K (10M)). The seven categories indicated by different colours identify voxels that show (a) significant co-activation and higher activation magnitude in WT, (b) significant co-activation and no significant inter-genotype difference in the activation magnitude, (c) significant co-activation and higher activation magnitude in HET, (d) significant co-deactivation and higher deactivation magnitude in WT, (e) significant co-deactivation and no significant inter-genotype difference in the activation magnitude, (f) significant co-deactivation and higher deactivation magnitude in HET, and (g) non-significant co-activation or co-deactivation during a CAP. Voxel-wise one-sample T-test (p < 0.01, Bonferroni corrected) and two-sample T-test (p < 0.05, FDR corrected) are performed across occurrences of a CAP within the concatenated genotypic image-series from all subjects to identify the voxels belonging to one of the 7 categories mentioned above and then divided among the 28 ROIs.

